# Optimal Therapy Scheduling Based on a Pair of Collaterally Sensitive Drugs

**DOI:** 10.1101/196824

**Authors:** Nara Yoon, Robert Vander Velde, Andriy Marusyk, Jacob G. Scott

## Abstract

Despite major strides in the treatment of cancer, the development of drug resistance remains a major hurdle. One strategy which has been proposed to address this is the sequential application of drug therapies where resistance to one drug induces sensitivity to another drug, a concept called collateral sensitivity. The optimal timing of drug switching in these situations, however, remains unknown.

To study this, we developed a dynamical model of sequential therapy on heterogeneous tumors comprised of resistant and sensitive cells. A pair of drugs (*DrugA, DrugB*) are utilized and are periodically switched during therapy. Assuming resistant cells to one drug are collaterally sensitive to the opposing drug, we classified cancer cells into two groups, *A_R_* and *B_R_*, each of which is a subpopulation of cells resistant to the indicated drug and concurrently sensitive to the other, and we subsequently explored the resulting population dynamics.

Specifically, based on a system of ordinary differential equations for *A_R_* and *B_R_*, we determined that the optimal treatment strategy consists of two stages: an initial stage in which a chosen effective drug is utilized until a specific time point, *T*, and a second stage in which drugs are switched repeatedly, during which each drug is used for a relative duration (i.e. *f*Δ*t*-long for *DrugA* and (1 – *f*) Δ*t*-long for *DrugB* with 0 ≤ *f* ≤ 1 and Δ*t* ≥ 0). We prove that the optimal duration of the initial stage, in which the first drug is administered, *T*, is shorter than the period in which it remains effective in decreasing the total population, contrary to current clinical intuition.

We further analyzed the relationship between population makeup, 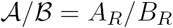, and the effect of each drug. We determine a critical ratio, which we term 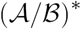, at which the two drugs are equally effective. As the first stage of the optimal strategy is applied, 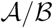 changes monotonically to 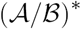 and then, during the second stage, remains at 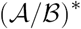 thereafter.

Beyond our analytic results, we explored an individual based stochastic model and presented the distribution of extinction times for the classes of solutions found. Taken together, our results suggest opportunities to improve therapy scheduling in clinical oncology.

## 1 Introduction

Drug resistance is observed in many patients after exposure to cancer therapy, and is a major hurdle in cancer therapy [1]. In most cases, treatment with appropriate chemo- or targeted therapy reliably reduces tumor burden upon initiation. However, in the majority of cases, resistance inevitably arises, and the disease relapses [2]. The observation of relapse is typically accomplished during surveillance through imaging, or in some cases a blood based marker [3, 4]. Disease recurrence is observed, at the earliest, when the disease burden reaches some threshold of detection, at which point the first line therapy is deemed to have failed and a second line drug is used to control the disease (see Figure 1 (a)). We argue herein that a redesign of treatment should start earlier than this time point, not only because the detection threshold is higher than the minimum disease burden, but also because the first drug could become less efficient as the duration of therapy reaches *T_max_*. In this research, we focus on the latter reason and figure out how much earlier we should switch drug in advance of *T_max_*, assuming that the former reason is less important (*t_DT_* – *t_o_*, ≈ *T_max_*).

**Figure 1:**
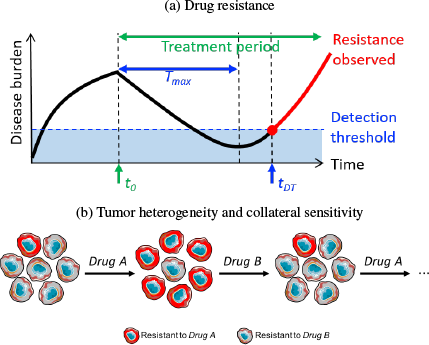
(a) General dynamical pattern of disease burden. It increases initially and then decreases as of the therapy starting point (*t*_0_), and eventually rebounds after the maximum period with positive therapy effect (*T_max_*). Relapse is found, at the earliest, when disease burden reaches detection threshold at *t_DT_*. (b) Change in composition of tumor cell population when a pair of collaterally sensitive drugs are given one after another.

While for many years it was assumed that tumors were simply collections of clonal cells, it is now accepted that tumor heterogeneity is the rule [5]. The simplest manifestation of this heterogeneity can be represented by considering the existence of both therapy resistant and sensitive cell types co-existing prior to therapy [6], with the future cellular composition shaped by the choice of drugs (illustrated in Figure 1 (b)). Beyond simple selection for resistant cells, cells can also become altered toward a resistant state during treatment, either by (i) genetic mutations [7, 8] or (ii) phenotypic plasticity and resulting epigenetic modifications [9, 10, 11].

To combat resistance, many strategies have been attempted, including multi-drug therapies targetting more than one cell-type at a time. While multi-drug therapy has enjoyed successes in many cancers, especially pediatric ones, the resulting combinations can often be very toxic. Further, recent work has suggested that the success of multi-drug therapy at the population level is likely overstated in individuals, given intra-patient heterogeneity [12]. Recently, researchers have sought specific sequential single drug applications that induce sensitivity, a concept is called collateral sensitivity [13, 14, 15, 16]. In some cases, several drugs used sequentially can complete a collateral sensitivity cycle [15, 14], and corresponding periodic drug sequence can be used in the prescription of long term therapies – though the continued efficacy of this cycle is not guaranteed [17]. In this research, we focus on a drug cycle comprised of just two drugs, each of which can be used as a targeted therapy against cells that have evolved resistance to the previous drug (illustrated in Figure 1 (b)).

The underlying dynamics of resistance development has previously been studied using cell populations consisting of treatment sensitive and resistant types, using either genotypic or phenotypic classifications [18]. Additionally, others have justified their choices of detailed cellular heterogeneities using: (i) stages in evolutionary structures [19, 20], (ii) phases of cell cycle [21, 22, 23, 24], or (iii) spatial distribution of irregular therapy effect [25, 26]. Among these, researchers (including [18, 22, 23, 27, 28]) have studied the effect of a pair of collaterally sensitive drugs as we propose here, using the Goldie-Coldman model or its variations [19, 28, 29, 30]. These models utilize a population structure consisting of four compartments, each of which represents a subpopulation that is either (i) sensitive to the both drugs, (ii) and (iii) resistant to one drug respectively, or (iv) resistant to both.

In this manuscript we propose a modeling approach which is the minimal model sufficient to study the effects of two populations of cells and two collaterally sensitive drugs. The model’s simplicity facilitates exact mathematical derivations of useful concepts and quantities, and illustrates several novel concepts relevant to adaptive therapy. The remainder of the manuscript is structured as follows. In Section 2, we outline the model and define terms. In Section 3 we present analysis of drug switch timing and duration. In Section 4 we relax several assumptions in our analytic model and study extinction times in a stochastic formulation, which agrees well with analysis in the mean field. In Section 5 we conclude and present work for future directions.

## 2 Modeling setup

### 2.1 Basic cell population dynamics under a single drug administration

Based on the sensitivity and resistance to a therapy, the cell population can be split into two groups. We refer to the population sizes of sensitive cells and resistant cells as *C_S_* and *C_R_* respectively, and then use the total cell population size, *C_P_* := *C_S_* + *C_R_*, to measure disease burden and drug effect. We account for three dynamical events in our model: proliferation of sensitive (*s*) and resistant cells (*r*), and transition between these cell types (*g*). Here, net proliferation rate represents combined birth and death rate, which can be positive if the birth rate is higher than the death rate or negative otherwise. It is reasonable to assume that, in the presence of drug, the sensitive cell population size declines (*s* < 0), resistant cell population size increases (*r* > 0), and that *g* > 0. Therefore, for the remainder of the work we consider only conditions in which *s* < 0, *r* > 0 and *g* > 0.

Figure 2 illustrates the population dynamics, and the system of ordinary differential equations that {*C_S_*, *C_R_*} obey. The solution of the system (1) is

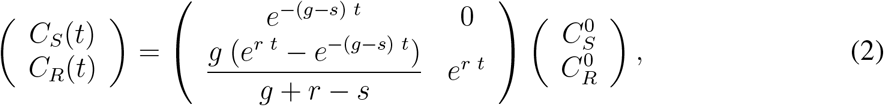

where 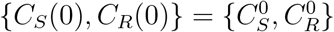. By (2), total population is

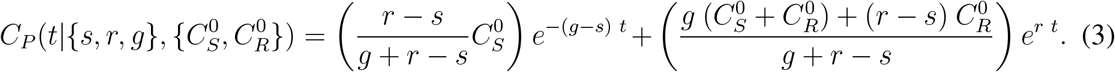

**Figure 2:**
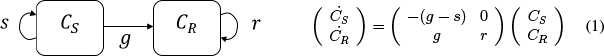
**Schematic of dynamics between sensitive cells population**, *C_S_*, **and resistant cells population**, *C_R_*, **(left panel) and the differential system of** {*C_S_,C_R_*} (**right panel**) with s–proliferation rate of sensitive cells, r–proliferation rate of resistant cells, (*g*–transition rate from *C_S_* tO *C_R_*

*C_P_*(*t*) is a positive function comprised of a linear combination of exponential growth (*e^r t^*) and exponential decay *e*^−(*g*–*s*)*t*^) with positive coefficients. Despite the limitations of simple exponential growth models [31], we feel it is a reasonable place to start, since the relapse of tumor size starts when it is much smaller than its carrying capacity which results in almost exponential growth.

*C_P_* has one and only one minimum point in {–∞, ∞}, after which *C_P_* increases monotonically. If 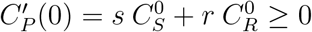, the drug is inefficient (*C_P_*(*t*) is increasing on *t* ≥ 0, see an example on Figure 3 (a)). Otherwise, if 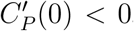, the drug is effective in reducing tumor burden at the beginning, although it will eventually regrow (due to drug resistance; see example in Figure 3 (b)).

**Figure 3:**
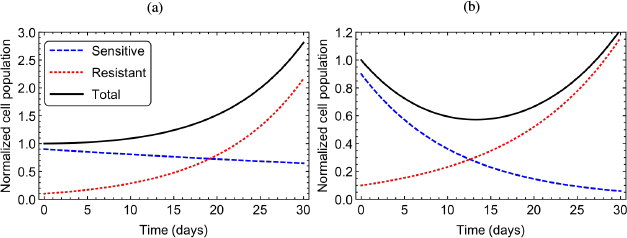
**Two representative population histories showing qualitatively different behaviors** depending on drug parameters with fixed initial population, 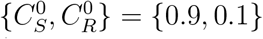. (a) increasing total population with {*s,r,g*} = {–0.01,0.1,0.001}; 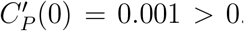. (b) rebounding total population with {*s, r, g*} = {–0.09,0.08, 0.001}; 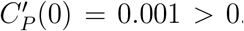.

### 2.2 Cell population dynamics with a pair of collateral sensitivity drugs

Here we describe the effect of sequential therapy with two drugs switched in turn, by extending the model for a single-drug administration (System (1)). Assuming that the drugs are collaterally sensitive to each other, cell population is classified into just two groups reacting to the two types of drugs in opposite ways. Depending on which drug is administered, cells in the two groups will have different proliferation rates and direction of cell-type transition (see Figure 4). That is, the population dynamics of the two groups follow a piecewise continuous differential system consisting of a series of the system (1), each of which is assigned to a time slot bounded by drug-switching times.

**Figure 4:**
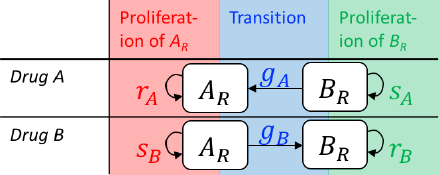
Dynamics of two cell subpopulations (*A_R_, B_R_*), which is opposite in direction under the present of collaterally sensitive drugs (*DrugA*, *DrugB*). *A_R_* is population of cells, resistant only to *DrugA*, and the *B_R_* population of cells, resistant only to DrugB in the presence of *DrugA* or *DrugB*. For each drug therapy, accounted cellular events are proliferations of sensitive and resistance cells ({*s*, *r*}, colored red and green) and drug-induced transitions from sensitive type to resistance type (*g* colored blue).

In summary, we assume that:

- There is a pair of collaterally sensitive drugs, *DrugA* and *DrugB*, which are characterized by their own model parameters: *p_A_* = {*s_A_,r_A_,g_A_*} and *p_B_* = {*s_B_,r_B_,g_B_*} respectively,
- A modeled tumor can be characterized entirely by two subpopulations, *A_R_* – resistant to *DrugA* and simultaneously sensitive to *DrugB*, and *B_R_* – resistant to *DrugB* and simultaneously sensitive to *DrugA*.
- Three factors determine the dynamical patterns, (i) drug parameters, {*p_A_, p_B_*}, (ii) the initial population sizes, {*A_R_*(0), *B_R_*(0)}, and (iii) the drug switching schedule.

An example of {*A_R_, B_R_, A_R_* + *B_R_*} histories is shown in Figure 5.

**Figure 5:**
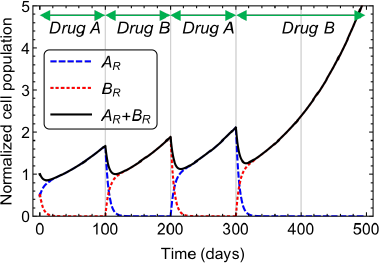
Representative plots demonstrating the dynamics of cell populations. Shown are population curves either resistant to *DrugA (A_R_*) or resistant to *DrugB (B_R_*), as well as the total population (*A_R_* + *B*_R_) during drug switches. Here, *p_A_ = p_B_ =* {–0.9, 0.08, 0.1}/*day* and {*A_R_*(0);*B_R_*(0)} = {0.5,0.5}.

## 3 Analysis of therapy scheduling

### 3.1 Drug-switch timing

To begin exploring the possible strategies of drug switching and timing within our model, we first tested an idea based on clinical intuition. As we discussed, the norm in the clinic is to change drugs when failure is *observed* either radiographically or through a bio-marker. We know, however, that the true failure occurs somewhat before this, yet at that time it is below the threshold of detection. To model drug switching at the point of ‘true failure’, the intuitive (yet unobservable) time point when the tumor population begins to rebound, we switch the drugs at the global minimum point of tumor size which we term *T_max_* (see Figure 1a), which was shown to exist uniquely in the previous section if and only if *C_R_*(0)/*C_S_*(0) < –*s/r*. The expression for *T_max_* derived from our model, is

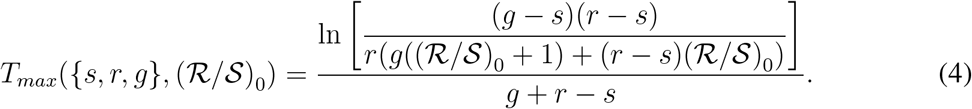

with (*R/S*)_0_ := *C_R_*(0)/*C_S_*(0). (See Appendix A.1 for this derivation.)

We see that the quantity *T_max_* depends only on (i) the parameters of the drug being administered, and (ii) the initial population makeup. In the *DrugA*-based therapy, it is 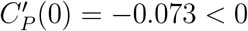, and in the *DrugB*-based therapy, it is 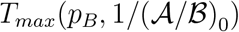, where 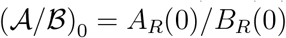.

In addition to *T_max_*, another important time point is *T_min_*, explained below. Since the rate of population decrease is almost zero around *T_max_*, with no switch (see the black curve of Figure 6), we seek to find a way to extend the high rate of population decrease by switching drugs before *T_max_*. To decide how much earlier to do so, we compared the derivative of *C_P_* under constant selective pressure (no switch) at an arbitrary time point, *t*_1_, and compared it to the right derivative of *C_P_* at *t*_1_ with the drug-switch assigned to *t*_1_.

**Figure 6:**
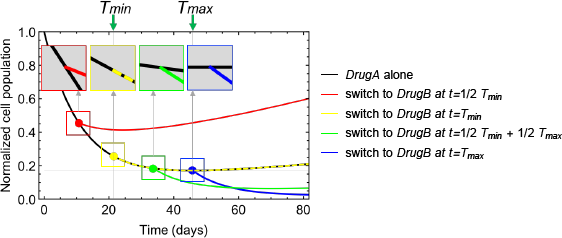
Comparison of total population curves with a one-time drug-switch from *DrugA* to *DrugB* at different time points. (i) at < *T_min_* (worse than without-switch; red curve), (ii) at *T_max_* (same as without-switch; yellow curve), (iii) between *T_min_* and *T_max_* (better than without-switch; green curve), and (iv) *T_max_* (better than without-switch; blue curve). Each color represents cell population size during and after a drug-switch using each switching strategy. The dashed yellow and black curve represents the overlap between the yellow and black curves. The tangent lines of the population curves at the chosen drug-switch time points are illustrated above. Parameters: *p_A_=p_B_* = {-0.9, 0.08, 0.001}/*day* and {*A_R_*(0), *B_R_(0)*} = {0.1, 0.9}.

**Figure 7:**
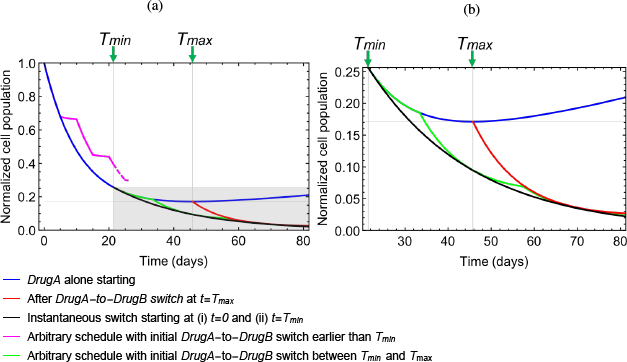
Total population curves with different therapy strategies with *p_A_ = p_B_ =* {–0.9, 0.08, 0.001}/*day* and {*A_R_(0), B_R_*(0)} = {0.1, 0.9} (a) full range of relative population (b) enlargement of the shaded areas on (a)

For example, if the first drug is *DrugA* and the follow-up drug is *DrugB* (illustrated in Figure 6), we compare

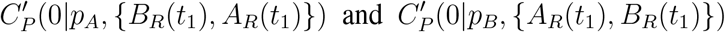

from (3). This comparison reveals that the two derivatives are equal iff *t*_1_ is a specific point (*T_min_*(see the yellow curve in Figure 6)) The derivative when the drugs are switched is lower (decreasing faster) iff *t*_1_ > *T_min_* (see the blue and green curves in Figure 6), and the derivative when the drugs are not switched is lower iff *t*_1_ < *T_min_* (see the red curve in Figure 6).

The general form of *T_min_* depends on the parameters of the “pre-switch” drug {*s*_1_, *r*_1_ *g*_1_} and for the “post-switch” drug {*s*_2_,*r*_2_}, as well as the initial population ratio between resistant cells and sensitive cells to the “pre-switch” drug, 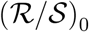 (See Appendix A.1 for details derivation). Here, the transition parameter in the second drug (*g*_2_), and the respective values of the two populations are unnecessary in the evaluation of *T_min_*, which is found to be

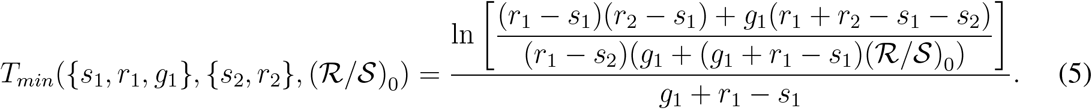

In the *DrugA*-to-*DrugB* switch, it is *T_min_(p_A_,p_B_*, (*A/B*)_0_), and in the *DrugB*-to-*DrugA* switch, it is *T_min_(p_B_,p_A_*, 1/(*A/B*)_0_)), where ((*A/B*)_0_) = *A_R_*(0)/*B_R_*(0).

It is important to note that the population curve with a single drug-switch after *T_min_* (and before *T_max_*, assuming that *T_min_* < *T_max_*) is not guaranteed to be lower than that of a single drug-switch switch at *T_max_* over the entire time range. As an example, as illustrated in Figure 6, the green curve relevant to the switch at (*T_min_* + *T_max_)/2* and the blue curve relevant to the switch at *T_max_* intersect at *t* ≈ 58 and the blue curve is lower after the time of this intersection. However, sequential drug switches starting between *T_min_* and *T_max_* create the possibility of finding a better drug schedule than the *T*_max_–based strategy. Figure 7 shows possible choices of follow up switches (green and black curves) which achieve better results than a *T*_max_–switch (red curves), unlike the drug-switches starting before *T_min_*, which remain less effective (magenta curve).

The optimal drug switching scheme will be discussed in detail in Section 4.2. The optimal scheduling for the example shown in Figure 5 starts by using the first drug until *T_min_* (blue curve for 0 < *t* ≤ *T_min_*) followed by a rapid exchange of the two drugs afterwards (black curve for *t* > *T_min_*)- Switching before *T_max_* that is, before the drug has had its full effect, goes somewhat against clinical intuition, and is therefore an opportunity for unrealized clinical improvement based on a rationally scheduled switch at *T_min_*. In order to realize this however, there are conditions about the order of *T_max_* and *T_min_* which must be satisfied. In particular:

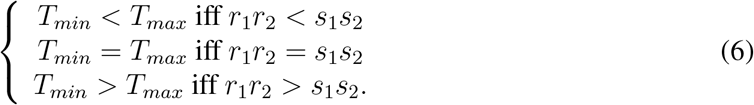

In our analysis and simulations, we will deal with the cases mostly satisfying *r*_1_*r*_2_ < *s*_1_*s*_2_, as otherwise the choice of drugs is not powerful to reduce the cell population (explained in detail in the next section and Figure 8).

**Figure 8:**
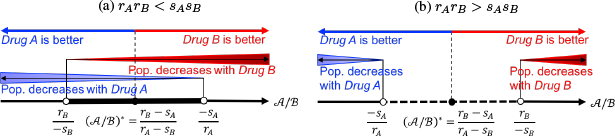
**Effect of** *DrugA* **and** *DrugB* **over the axis of** 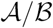. The two drugs have the same effect when *A/B* = (*A/B)\**, and have no effect when 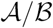 = *–s_A_/r_A_* (in the case of *Drug A) or A/B = –r_B_/s_B_* (in the case of *DrugB*). The drug effect increases as *A/B* gets farther from the no-effect level in the direction a smaller resistant subpopulation. Depending on a condition, there exists (Panel a) or does not exist (Panel b) a range of 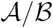 in which both drugs have positive effects.

This window of opportunity, where the clinical gains could be made, which we will term *T_gap_*, is the difference between *T_min_* and *T_max_*. This relationship allows us to compare *T_min_* and *T_max_* using different parameters.

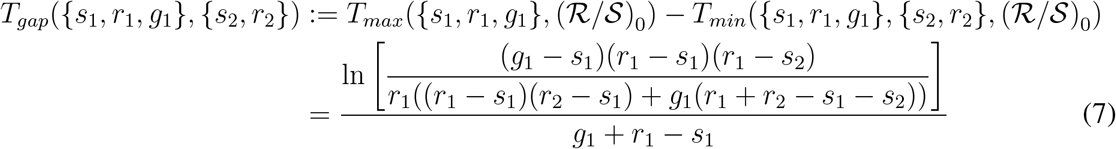

We analyze sensitivity of *T_gap_* over a reasonable space of non-dimentionalized drug parameters in Appendix B. As expected, as the proliferation rates under the second drug increases (*r*_2_ ↑ and/or *s*_2_↑), the optimal time to switch to the second drug is delayed (*T_min_* ↑ and *T_gap_* ↓). As *r*_1_ increases, both *T_min_* and *T_max_* decrease. However, *T_max_* decreases more than *T_min_* does, so overall *T_gap_* decreases, *s_1_* and *T_gap_* do not have a monotonic relationship. As *s*_1_ increases, *T_gap_* increases for a while (when *s*_1_ is relatively low), and then decreases afterward (when *s*_1_ is relatively high).

### 3.2 Population makeup and drug effect

In the previous section, the derived time points (*T_min_*, *T_max_*) are dependent on the initial population makeup (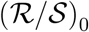) from Equations (4)-(5), but not on explicit size of the total population or subpopulations. This makes sense, since absolute population size plays a role by scaling overall behavior of populations 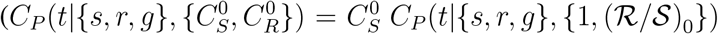 from (2)), and *T_min_* and *T_max_* are both defined by derivatives at the time points (i.e., *C_P_*(*T_max_*) = 0, and from (5)). In this section, we seek to clarify the relationships between population makeup and therapy effects defined using 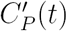, and roles of *T_min_* and *T_max_* in these relationships. We first define functions of the ratio between the two cell subpopulations:

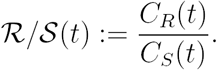

We further define functions measuring drug effectiveness as the relative rate of population change depending only on 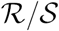 and drug parameters:

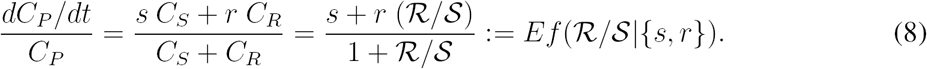

In the case where we classify cells as *A_R_* and *B_R_*, we similarly define their population makeup as:

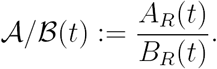

Then 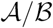 at *T_min_*, using a *DrugA*-to-*DrugB* switch 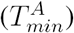, and *A/B*, using a *DrugB*-to-*DrugA* switch 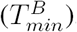, are equivalent:

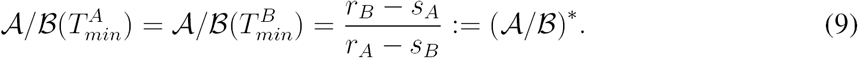

At *T_max_* with *DrugA* 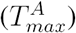, and with *DrugB* 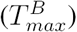, we have

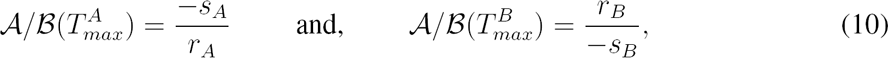

and further, as *s* < 0 and *r* > 0, values of 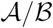 are all positive. We give a more thorough description of (9) and (10) in Appendix A.1.

The effects of *DrugA* (specified by *p_A_*) and *DrugB* (specified by *p_B_*), both defined by (8), are equivalent at *T_min_*, that is 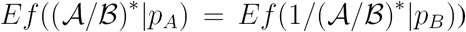. The effect of *DrugA* is larger if 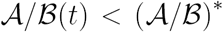, since the *DrugA* resistant cell population is relatively smaller than the population of the other cell type, otherwise, *DrugB* has a more beneficial effect. When *t* = 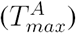 and therefore when 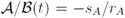, *DrugA* has no effect on population reduction (i.e. *Ef(–(s_A_/r_A_|p_A_*) = 0). If 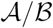 is getting smaller, *DrugA* becomes effective. Furthermore, the smaller 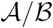 is, the better the effect *DrugA* has. Similarly the effect of Drug B is zero when *t* = 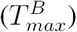 and 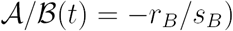 and increases as 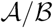 increases above it (see Figure 8).

The population makeup changes in the opposite direction as *DrugA* (or *DrugB*) therapy continues, 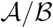 therefore continues to increase (or decrease). Therefore, if *DrugA* (or *DrugB*) is given too long, it goes through a period of no or almost no effect around 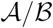 = –*sA/rA* (or around 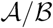 = *–r_B_/s_B_*), but once the drug is switched after that, there will be a higher therapy effect with *DrugB* (or with *DrugA*). These two opposite aspects are balanced by switching the drug when the population makeup reaches 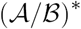, which is applied to the optimal therapy regimen described in the next section.

Depending on condition (6), the order of the three population makeups at *T_min_*, 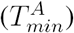 and 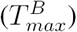 changes. In particular, if *r_A_r_B_* < *s_A_s_B_*, there exists an interval (–*r_B_/s_B_,–s_A_/r_A_*) in 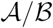 in which both drugs are effective in decreasing the population size. Otherwise, if *r_A_r_B_* < *s_A_s_B_*, no drug is effective when 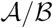 ∈ –*s_A_/r_A_*. –*r_B_/s*_B_). These results are illustrated in Figure 8.

### 3.3 Optimal scheduling and its clinical implementation

In this section, we describe a drug-switching schedule design to achieve the best effect possible with a pair of collaterally sensitive drugs. The area under the curve of the total population simulated under an assigned treatment strategy is utilized to measure the aggregate effect of the strategy. The smaller the area, the better the corresponding strategy. The numerically determined optimal strategy consists of two stages:

- **Stage 1:** Treat with first drug until reaching the population makeup where the effects of each drug are balanced (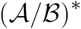), that is until the *T_min_* of the first drug.
- **Stage 2:** Begin switching drugs with a specific temporal ratio (represented by *k* or *k*’, see Figure 9) determining the difference in the treatment duration of each drug, and switching frequently (represented by Δ*t* ≈ 0). Both conditions are used to keep 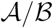 close to constant near 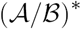.

**Figure 9:**
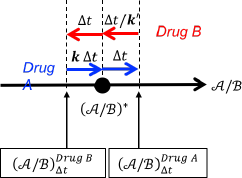
Schematic of the relationship between therapy duration (*Δt, k Δt*, or Δ*t/k’*) and the change in 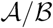 around 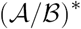. *Δt* represents an arbitrary time interval (ideally short, *Δt* ≈ 0) and *k* represents a specific quantity corresponding to *Δt* and the model parameters in *DrugA* and *DrugB*.

We represent the relative durations of *DrugA* compared to the duration of *DrugB* in Stage by *k* and *k’*. The explicit formulation of *k* can be derived from the solution of the differential equations (2). To do so we (i) evaluate the level of 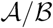 after *Δt* time has passed during *DrugA* therapy, when starting with 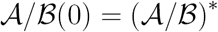, that is 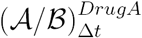, and then (ii) by measuring the time period taken to regain (*A/B)\** from 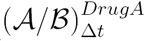 through therapy with *DrugB*, denoted by *At’*, and finally (iii) taking the ratio between the two therapy periods, which is *k* := *Δt/Δt’, k* depends on the frequency of drug switching and model parameters:

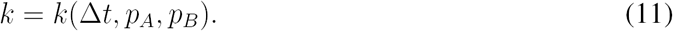

This *k* is consistent with *k’* = *k’*(Δ*t*, *p_A_, p_B_*), which is the ratio similarly evaluated with *DrugB* as the first therapy and *DrugA* as the follow-up therapy, in the optimal case of instantaneous switching:

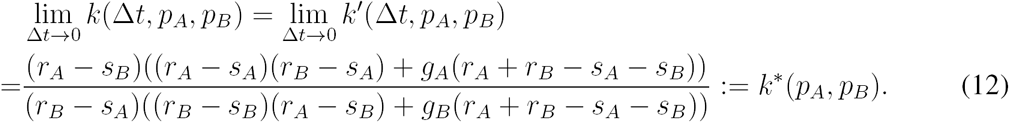

For a more detailed derivation of *k\**, see Appendix A.1. We further studied how sensitive *k\** (or *f*\* = *k*/(* 1 + *k*)*) is over a reasonable range of non-dimentionalized *{p_A_,p_B_}* (see Appendix B for details). *k\** (or *f*\*) increases, as *r_A_* and/or *s_B_* decreases as *s_A_* and/or *r_B_* increases.

Figure 10 shows examples of population curves with the optimal strategy (*T_min_* switch) and one non-optimal strategy (*T_max_* switch) using the same choice of parameters/conditions. Visual comparison of total population curves (Figure 10 (a)) reveals that the predicted optimal strategy outperforms the intuitive strategy. To quantitatively compare the efficacy of each strategy, we can use area between the two population curves. This area is:

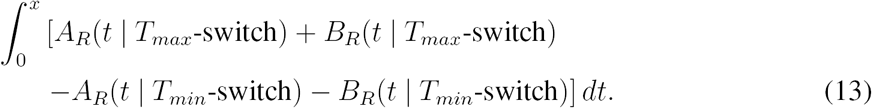

**Figure 10:**
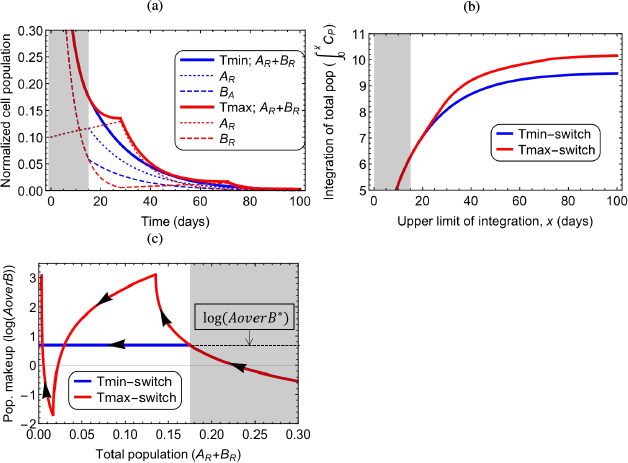
Comparison between dynamical trajectories using the optimal (*T_min_* switch; blue curves) and an example of non-optimal (*T_max_* switch; red curves) therapeutic strategies, in terms of (a) time histories of *A_R_, B_R_* and *A_R_* + *B_R_*, (b) integration of total population from *t =* 0 to varying upper limit (x-axis), and (c) dynamical changes in the total population and population makeup. On all panels, Stage 1 is shown in gray and Stage 2 is shown in white. Parameters/conditions are: {s_A_,s_B_} = {–0.18, –0.09}/day, {*r_A_,r_B_}* = {0.008,0.016}/day, {*g_A_,g_B_*} = {0.00075, 0.00125}/day and 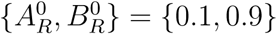.

With a choice of upper limit large enough to include most treatment schedules, *x* = 100 (days), we used sensitivity analysis of the integral (13) (See Appendix B for the details). The advantage of the optimal treatment strategy is demonstrated by the lower population sizes in all cases. And the evaluations of the areas under the population curves from *t* = 0 to a range at the upper limit of integration (Figure 10 (b)) confirms the superior effect of the optimal strategy over time. Figure 10 (c) shows the typical pattern of 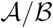 in the optimal therapy compared to the other, which is monotonically changing toward 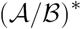 in the first stage and constant in the second stage.

While our theory predicts optimality with “instantaneous drug switching”, we realize this is not clinically feasible. Therefore, the instantaneous drug switching in Stage 2 could be approximated by a high frequency switching stratgey with *Δt* ≳ along with the corresponding *k(Δt*) from (11), or *k\** (12) independent from *Δt*. As expected, the smaller *Δt* is chosen, the closer the population follows the ideal case with *Δt* = 0 (see Appendix C for the details), but improvements can still be made over non-strategic switching, if the temporal ratio is followed.

We have proved that the effect of instantaneous drug switching, with an arbitrary ratio in duration between two drugs (*k*), is consistent with the effect of a mixed drug with a relative dosage ratio, which is also *k* (Theorem A.8 in Appendix A.2). The theorem is used in the derivation of a differential system/solution of the optimal strategy (Theorem A.11 in Appendix A.3). According to these results, in Stage 2 of optimal regimen, all types of populations, *A_R_, B_R_* and *A_R_* + *B_R_*, change with the same constant proliferation rate:

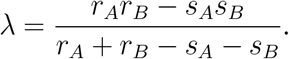

While not clinical proof, these theoretical results suggest a method of application of two drugs in sequence, which would approximate multi-drug therapy in efficacy, but which could be free of the increase in side effects from the combination.

## 4 Studying extinction time with a stochastic formulation

In the previous sections we utilized an entirely deterministic model of heterogeneous tumor growth. Cancers, however, are not deterministic, and without stochasticity in our system we could not model an important part of cancer treatment: extinction. We therefore constructed a simple individual based model using a Gillespie algorithm [32] to study this critical aspect of therapy that is not limited by the assumptions we were required to make for purposes of analytic tractability.

Our stochastic model depends not only on net proliferation rates (*s*, *r*, see Equation (1)) but also on the combination of birth rates (*b_S_, b_R_*) and death rates (*d_S_, d_R_*) where *s = b_S_ – d_S_* and r = *b_R_ – d_R_*. These five parameters (*b_s_, b_r_, d_s_, d_r_, g*) govern the probabilities of events occurring. The time at which one of these events occurs is determined by an exponential probability distribution, and we represent the algorithm as pseudo-code thus:

**(Step 1)** Initialize 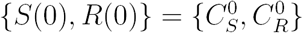.

**(Step 2)**

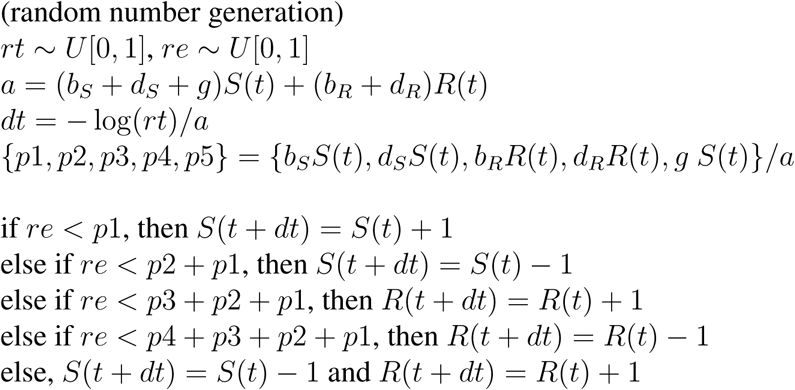

**(Step 3)** *t* ⟵ *t* + *dt* and repeat **(Step 2)** until a set time has passed or extinction has occurred.

We expanded the stochastic process for a single drug to treatment with two drugs being switched in turn, as in our ODE system (See Appendix D, for the details of the computational code). Figure 11 (a) shows the consistency between the mean field behavior of the stochastic model and the ODE system.

**Figure 11.**
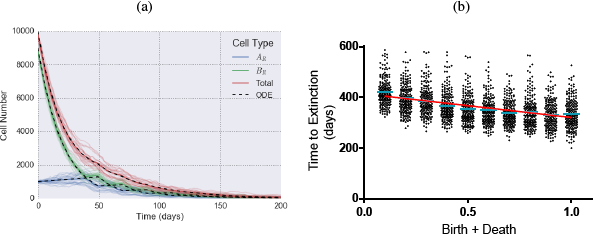
(a) Comparison between the stochastic process and the ODE model. The mean (thick curves) of multiple stochastic simulations (thin curves) are compared to the ODE solution (dashed curves). Parameters are *{s_A_,r_A_, g_A_|s_B_,r_B_,g_B_*|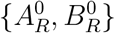 = {–0.05,0.005,0.– 0.05,0.005,0.000111000,9000}, birth rate + death rate (*I_stoch_*) = 1.0. (b) Relationship between birth-death combinations (*I_stoch_*; 0.1 to 1.0 with intervals of 0.1) and simulated extinction time in replicates with the same parameters and initial condition with (a). Regression (red line) is *y* = –93.68x + *α* (slope has p<0.and *r*^2^ = 0.1726). Cyan lines show mean values.

Despite the generally similar patterns of population curves simulated with same {*s, r, g*}-type parameters and initial conditions, we observe differences in terms of elimination time if birth/death combinations are different. To quantify these differences we directly studied the elimination times (defined as the distribution of times to the absorbing state of total population = 0) simulated with different combinations of birth/death rates, with a choice of fixed proliferation rates (as well as other fixed transition rates and initial condition). We defined an index to represent different levels of birth and death rate combinations:

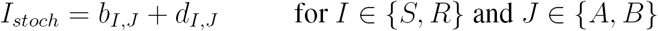

where *I* indicates a type of sensitivity or resistance and *J* does a type of drug. Given a specific net proliferation rate (*b_I,J_* – *d_I,J_*), the larger the index, the larger both birth (*b_I,J_*) and death (*d_I,J_*) rates are.

Increased *I_stoch_* result in larger fluctuations, these fluctuations then increase the probability of reaching the absorbing state which is extinction (tumor cure). The relationship between *I_stoch_* and extinction time is shown in Figure 11 (b). The relationship is approximated by a linear model with slope, −93.68 (days^2^), p-value of the slope, *p <* 0.05, and squared residual of regression, *r*^2^ = 0.1726.

## 5 Conclusions and discussion

The emergence of resistance to the best current cancer therapies is an almost universal clinical problem, and the solution to this represents one of the greatest unmet needs in oncology. While much effort has been put into novel drug discovery to combat this, there is also a growing interest in determining the optimal sequences, or cycles of drugs that promote collateral sensitivity. To study this second paradigm, we proposed a simple dynamical systems model of tumor evolution in a heterogeneous tumor composed of two cell phenotypes. While in reality, cell phenotype can be defined in many ways, here we completely describe it by considering only sensitivity (or resistance) to a pair of collaterally sensitive drugs, which is encoded in their differential growth rates in specific conditions. While the resulting mathematical model conveys only simple, but essential, features of cell population dynamics, it does yield analytical solutions that more complex models cannot.

Our original motivation was to consider more complicated sequences, or cycles of drug therapy, however, the model presented herein is difficult to apply for an expanded system of more than two drugs. On the other hand, the cell classification used by others [18, 19, 28, 29, 33] considers sensitivity and resistance independently, or even specifically to a given, abstracted, genotype [34, 35]. Therefore, in the case of 2 drugs, there are 2^2^ = 4 groups, (i) sensitive to both drugs, (ii) and (iii) resistant to only one drug, and (iv) resistant to both drugs. This formulation could be expanded and applied to more than two drugs [18, 33]. Also, in other earlier researches, cell populations are divided by more specific criteria for the choices of cancers and drugs (e.g., level of protein expression, enzyme inhibitors, or growth factors [10, 11, 8]). We will consider both of the general and specific approaches of population classification in future work.

The simplicity of our exponential growth/decay model arises from the assumption of a constant growth rate. Use of exponential growth is likely not overly inappropriate, as we are most interested in the development of resistance – and resistance is typically thought to begin when the tumor burden is much smaller than the carrying capacity. However, the assumption might have oversimplified patterns of cell growth, which is assumed to be non-exponential by others (e.g. logistic growth [31, 36, 37]), due to the limited space and resources of the human body for tumor growth, as well as increasing levels of resistance (increasing growth rates) in the face of continued selective pressure [38]. We will consider the concept of changing growth rates in terms of time and population density, and explore its effect on our analytical results (such as *T_gap_*, 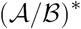, *k\** and etc.) in future work.

We provided a strategy for drug-switching which yields the best possible effect in this model system, i.e. the fastest decrease in cell population. The strategy is defined explicitly in terms of parameters determined by the drugs that are used, therefore the applicability of our model relies on the availability of drug parameters. Drug parameters for several drugs are known based on *in vitro* experiment or clinical studies [39, 40]. However, these parameters are not available for all drugs, and even the usefulness of *in vitro* results may change from one patient to the next. Because of this, we propose focusing our future work on learning to parameterize models of this type from individual patient response data. Examples of parameterizing patient response from imaging [41] as well as blood based markers [42] already exist, suggesting this is a reasonable goal in the near future.

In our optimized treatment regimen we must first apply *DrugA* (if *DrugA* is better at the initial time, i.e., 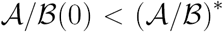, see Figure 8). Surprisingly the ideal treatment course switches to *DrugB* while *DrugA* is still effective at reducing the total population. Since treatment should ideally switch before the tumor relapses our study justifies the search for techniques that either identify or predict resistance mechanisms early. Our study also argues against the opposite extreme, wherein resistant cells are targeted at the beginning of treatment. The preponderance of cells sensitive to the standard of care makes this treatment initially ideal, and does not preclude eventual success in our model. Further, the rapid tumor size reduction, associated with targeting the larger sensitive population first, could be clinically meaningful.

Our stochastic model allowed us to explore the contributions of cell birth and death separately, as opposed to the ODE which could only consider the net growth rate. These parameters can be altered in cancer since cancer treatments have various cytostatic and cytotoxic effects, and therefore different treatments can have different effects on death and birth. In our model, increasing the total birth and death rate (as opposed to the net growth rate) caused, on average, extinction earlier in time (Figure 11 (b). This can be explained by the fact that extinction is the only absorbing state in our model, and therefore higher death rates determine when extinction occurs, even when birth rates are also higher. Our stochastic model therefore suggests that highly cytotoxic drugs (even those with correspondingly minimal cytostatic effects) are more effective at eliminating tumors, at least when the tumor population is small.

In summary, we have presented a simple model of a heterogeneous, two phenotype tumor, with evolution occurring between resistant and sensitive states. We derive exact analytic solutions for tumor response in temporally changing drug conditions and find an optimal regimen which involves drug switching after a specific, critical time point which occurs before resistance would normally be clinically evident. While our model is highly simplified, we have identified several opportunities to improve our understanding and treatment of drug resistance, and also future opportunities for new modeling endeavors.

## Appendix A Derivations of explicit expressions

### A.1 Details of Equations (4), (5), (6), (7), (9), (10) and (12)

1. *T*_*max*_: Equation (4)

*T_max_* is a minimum point of *C_p_(t*) (from (3)). Therefore,

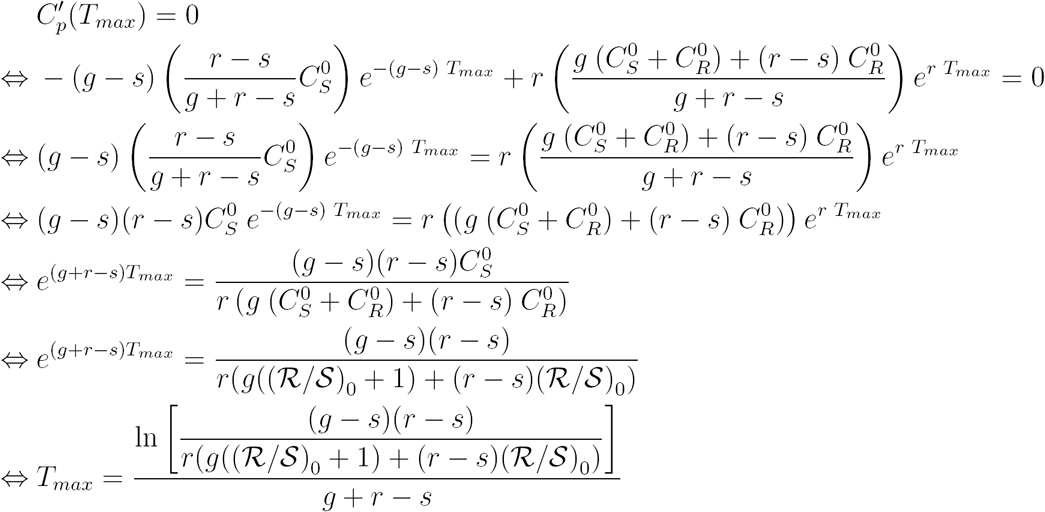

2. *T*_min_: Equation (5)

Let us consider the case of drug switch with *DrugA* being the “pre-switch” drug and *DrugB* being the “post-switch” drug. If, at a specific time point *t*_1_, cell population is decreasing faster 5u by continuing *DrugA*-therapy than by changing drug to *DrugB*,

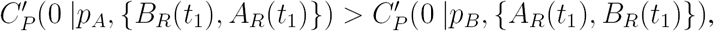

from Equation (3) where 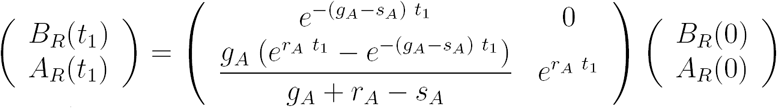 evaluated from Equation (2). Then,

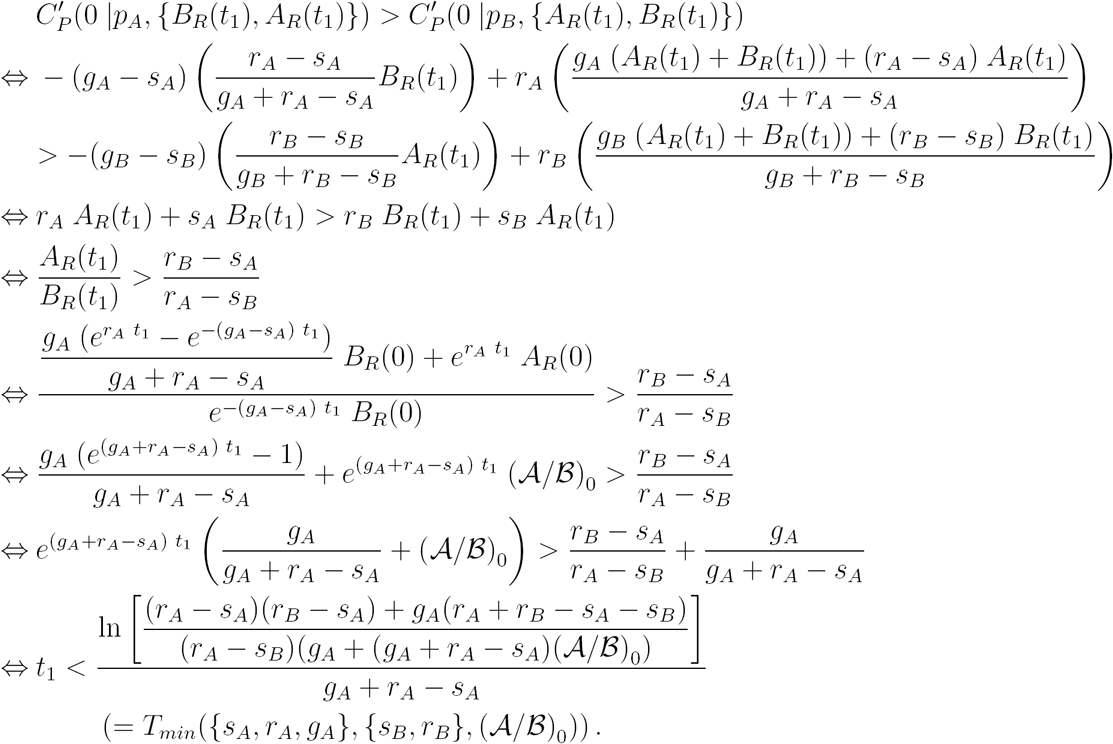

Similarly,

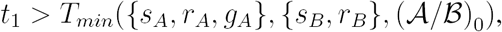

iff the population is dropping faster using *DrugB* than by continuing to use *DrugA*, and

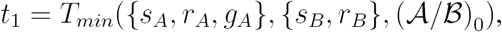

iff the population is dropping at an equal rate with either drug.

The general form of *T_min_* is

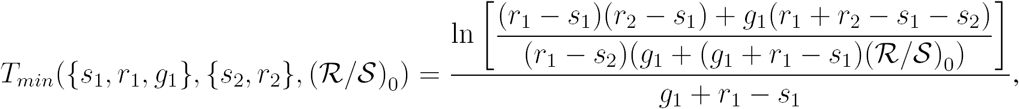

where the parameters of “pre-switch” and “post-switch” drugs are {*s*_1_, *r*_1_, *g*_1_ and {*s*_2_, *r*_2_. *g*_2_} respectively, and initial population makeup, 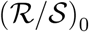, is the resistant cell population divided by the sensitive cell population for the “pre-switch” drug.

3. *T_gap_:* Equation (6) – (7)

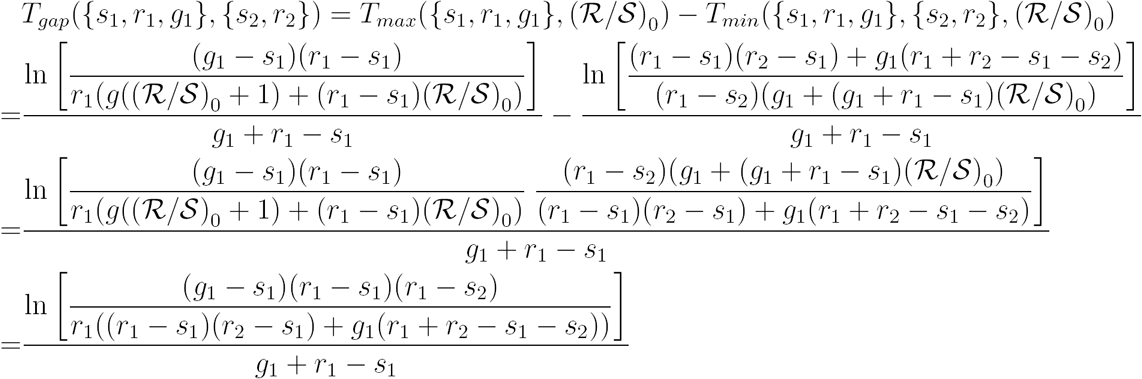

And,

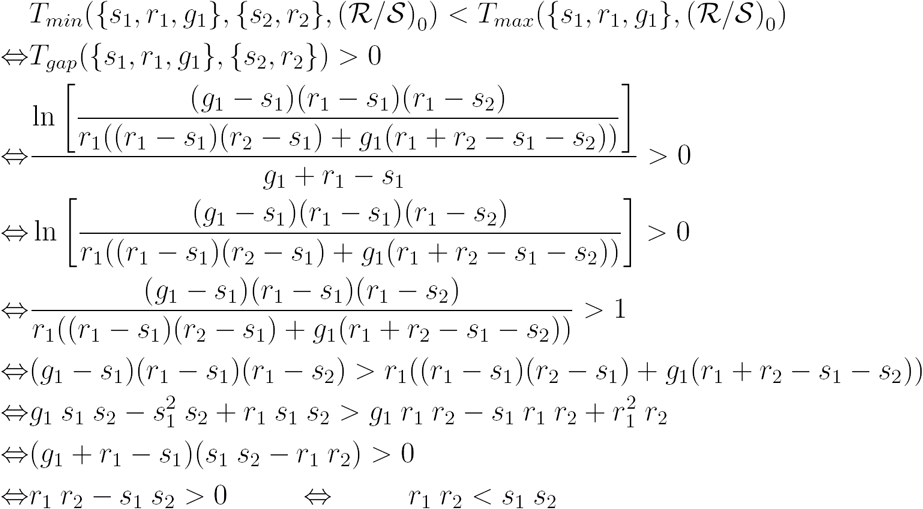

Similarly *T_gap_* = 0 iff *r*_1_ *r*_2_ = *s*_1_ *s*_2_, and *T_gap_* < 0 iff *r*_1_ *r*_2_ > *s*_1_ *s*_2_.

4. 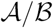 at *T_max_* and *T_min_*: Equation (9) – (10).

It is clear that

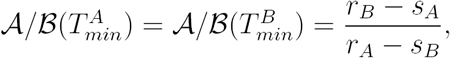

and

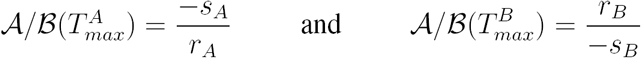

by the expressions of *A_R_(t), B_R_(t), T_max_* and *T_min_* from Equations (2), (5) and (4).

Otherwise, it can be proved more simply using the concept of *T_min_* and *T_max_*. Since 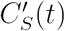 + 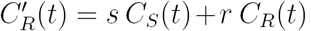, from the differential system (1), the derivatives of *A_R_(t) + B_R_(t*) are *s_A_ B_R_(t*) + *r_A_ A_R_(t*) and *s_B_ A_R_(t*) + *r_B_ B_R_(t*) under *DrugA* and *DrugB* respectively. At *T_min_* (whether it is 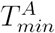 or 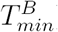) the derivatives of total populations are equivalent either under *DrugA* or under *DrugB*. Then,

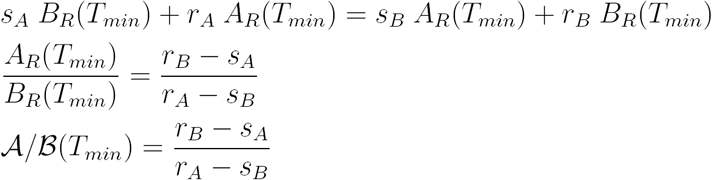

Therefore,

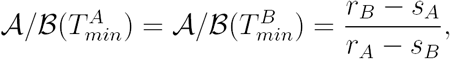

Under *DrugA* at 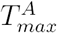. Therefore,

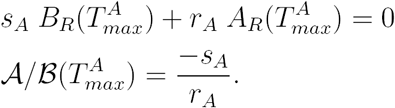

Similarly, 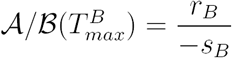.

5. *k*:* Equation (12) The sizes of the subpopulations after Δ*t*-long therapy with *DrugA* started from initial population makeup of 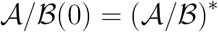 are

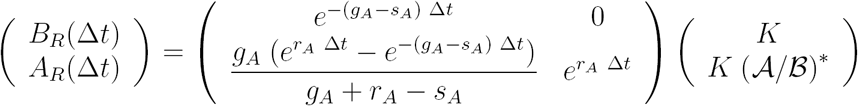

derived from Equation (2), with some constant *K* scaling population size. Then the population makeup at the Δ*t* and its derivative in terms of Δ*t* are

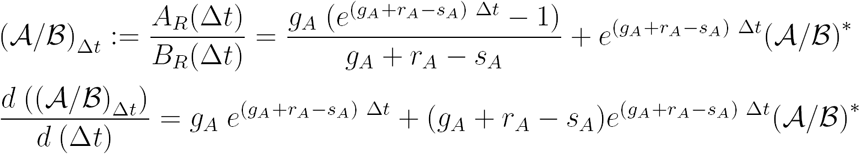

The time taken from *t* = Δ*t* to reach back to the time of 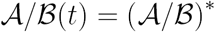 given *DrugB* is

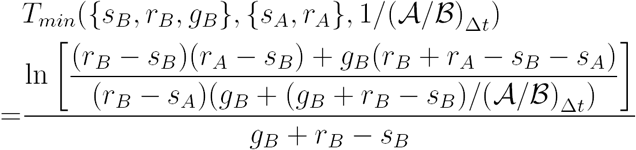

from Equation (5).

Then the relative ratio between the periods of *DrugA* and *DrugB, k’*, illustrated in Figure 9, and its limit, *k\**, can be derived using:

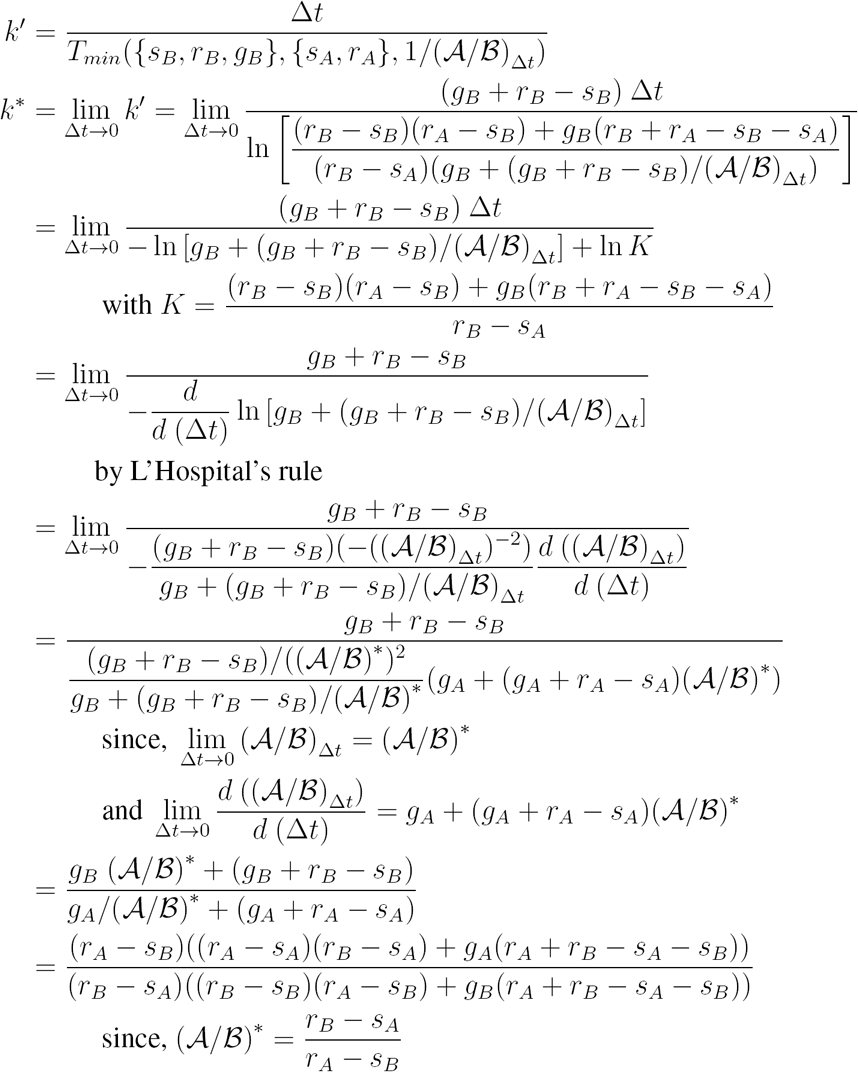

### A.2 Differential system of 552 instantaneous drug switch

The goal of this section is to derive the simple differential equations of *V* = {*A_R_, B_R_*} under instantaneous drug switch (Theorem A.8). For the sake of convenience, we want to use matrix operations and equations based on the vectors and matrices defined below.

#### Definition

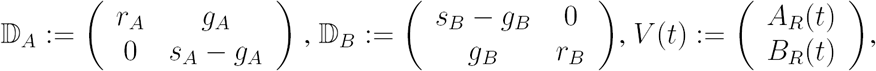

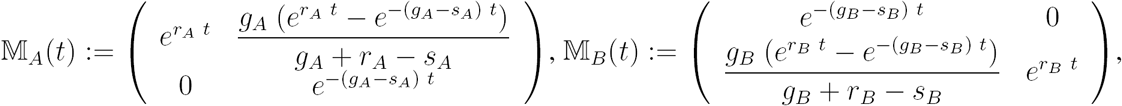

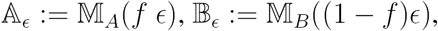

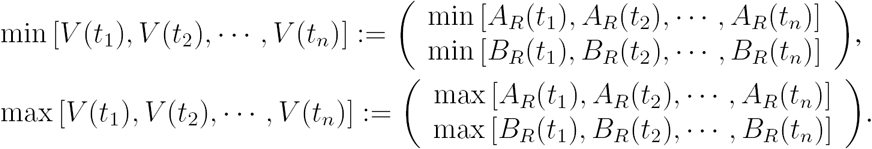

**Proposition A.1.** *Using Drug A therapy:*

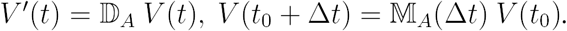

*Using Drug B therapy:*

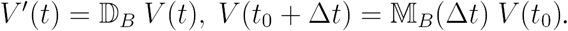

**Proposition A.2.** *Both A_R_ and B_R_ are monotonic functions under either therapy. In the presence of Drug A, A_R_ is increasing, and B_R_ is decreasing. And, in the presence of Drug B, A_R_ is decreasing, and B_R_ is increasing.*

**Proposition A.3.** 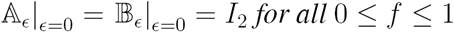

**Proposition A.4.** 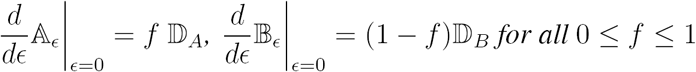

**Lemma A.5.** 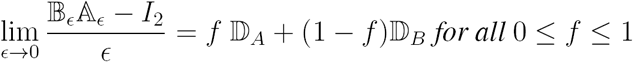

Proof.

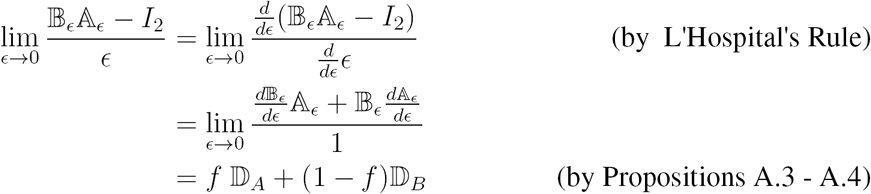

**Lemma A.6.** 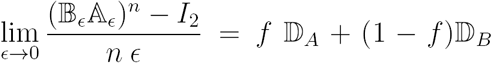 *for any positive integer, n, and for all* 0 ≤ f ≤ 1

*Proof.* Let 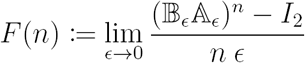 and 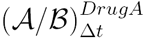. Then, we need to prove that *F(n)* = *L* for *n* = 1,2,3,…

If *n* =1,

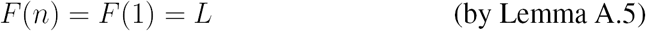

Otherwise, if *n* ≥ 2 and *F(m)* = *L* for all 1 ≤ m ≤ *n* – 1,

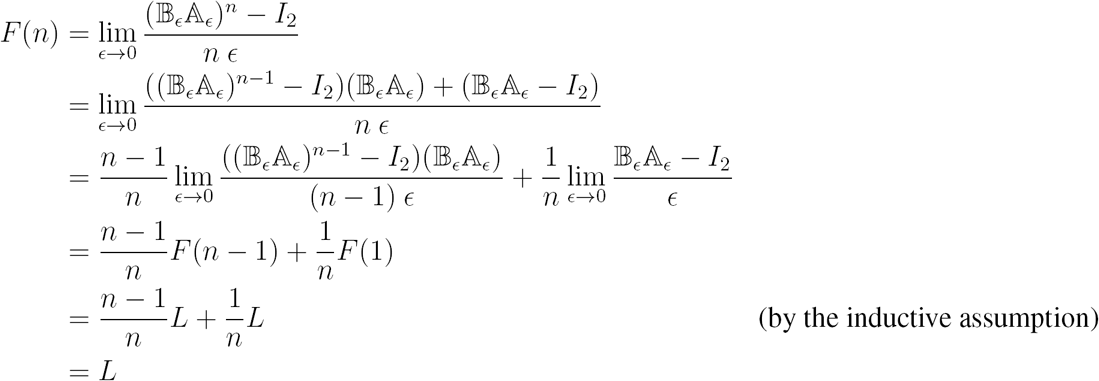

Therefore, proved.

**Lemma A.7.** 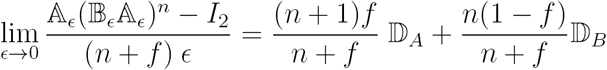 *for any positive integer, n, and for all* 0 ≤ *f* ≤ 1

*Proof.* Using mathematical induction, if *n* = 1,

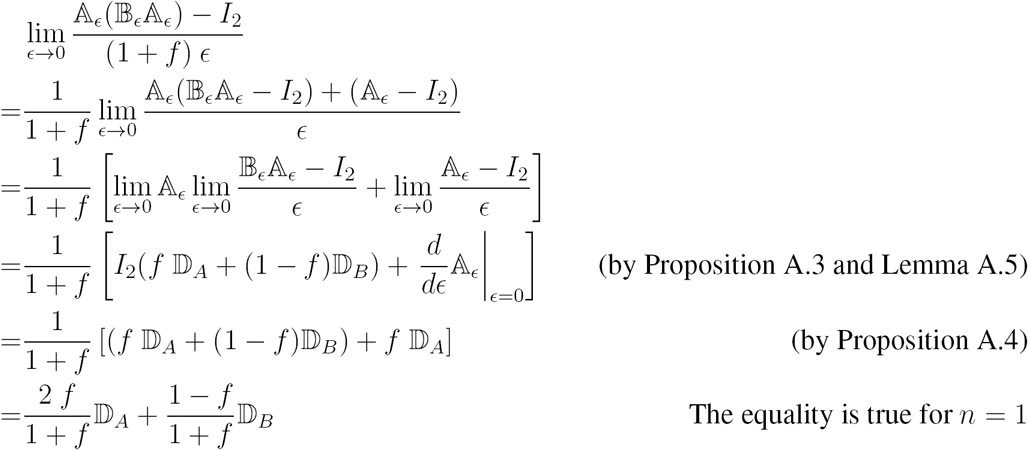

If n ≥ 2, and the equality works for all integers 1 ≤ *m* ≤ n – 1,

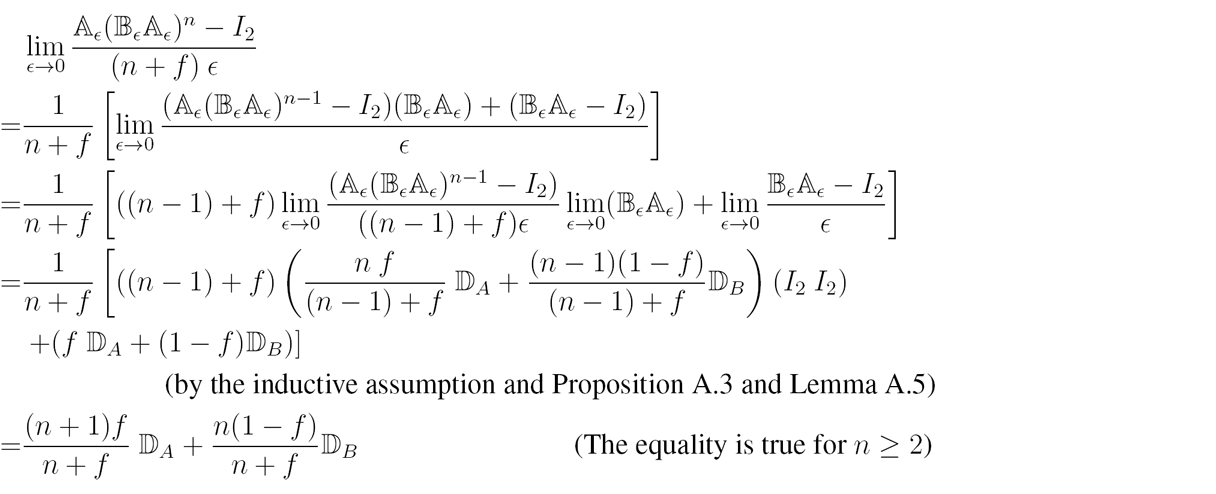

Therefore, proved.

**Theorem A.8.** *If Drug A and Drug B are prescribed in turn with a relative intensity of f and 1 – f, and are switched instantaneously, V obeys*

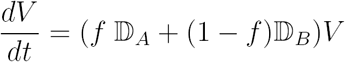

*Proof*. For any time point *t*_0_, let us define *V_ϵ_(t)* as a vector-valued function of *A_R_(t)* and *B_R_(t)* describing the cell population dynamics under a periodic therapy starting at *t_0_* with *DrugA* assigned at *t*_0_ + *m ϵ ≤ t < t*_0_ + (*m + f)ϵ* and *DrugB* at *t*_0_ + *m ϵ ≤ t < t*_0_ + (*m + 1)ϵ* for *m* = 0,1,2, 3,…. Then, by Proposition A.1 and the definitions of 픸 and 픹,

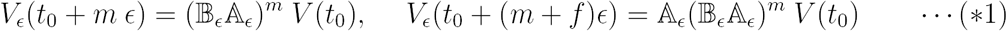

where 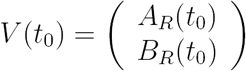. And, V_0_(*t*) represents instantaneous drug switching.

For any *Δt >* 0 and any positive integer *n*, there exists *ϵ =ϵ(n, Δt*) such that

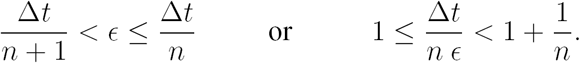

Then by the squeeze theorem,

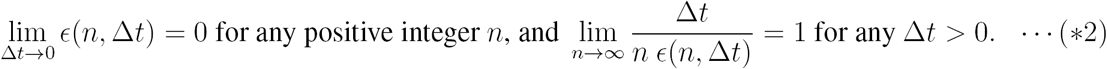

For such *Δt, n* and *ϵ(n, Δt), V_ϵ_(t_0_* + *Δt*) is bounded, since local extrema can occur only when drugs are switched by Proposition A.2. That is,

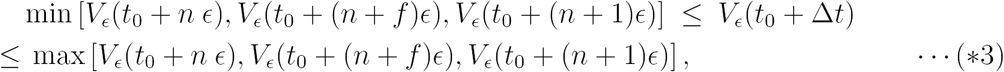

Also,

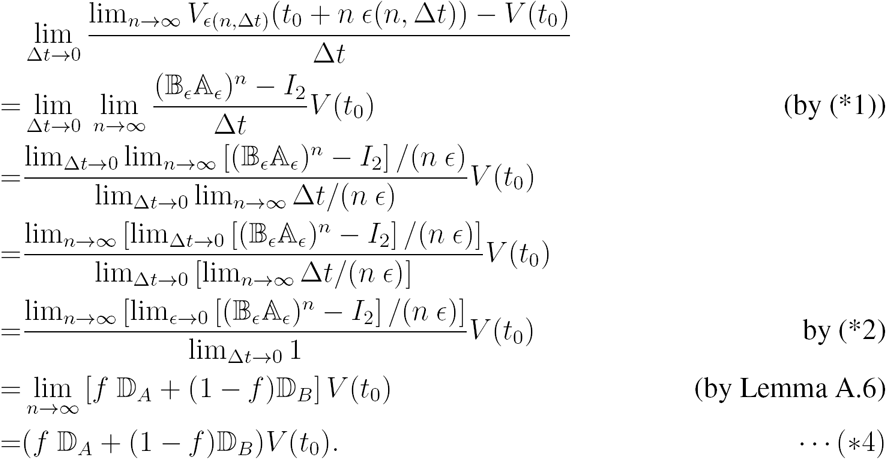

And,

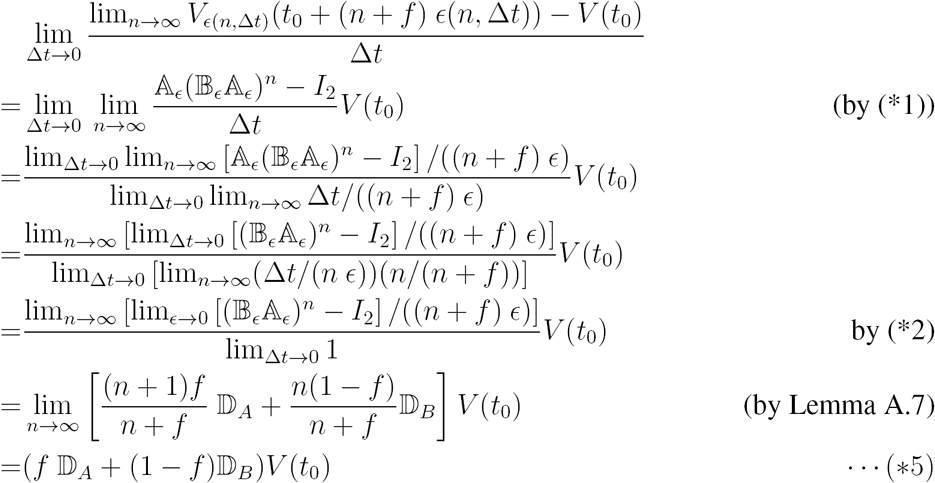

Similar to (*4),

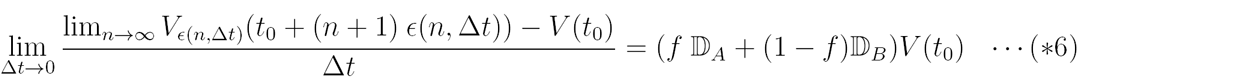

By (*4) – (*6),

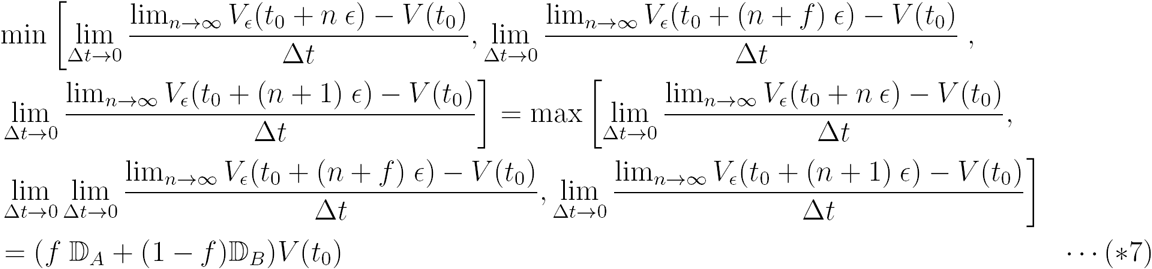

Then, by (*3), (*7) and 585 the squeeze theorem,

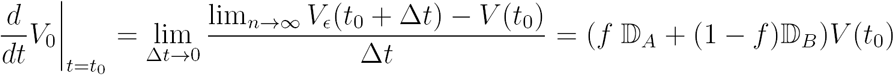

Therefore,

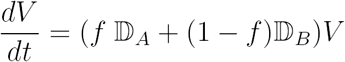

### A.3 Population dynamics with the optimal regimen

In this section, we want to write the differential equations of *V* = {*A_R_,B_R_*} under the optimal control strategy described in Section 3.3. Based on Appendix A.2 and a couple of lemma/theorem, we will reach to a concise form of a differential system described at Theorem A.11.

Lemma A.9. 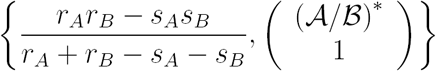 is an eigen pair of 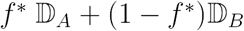 with (A/B)* and 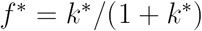 defined by Equations (9) and (12).

*Proof.* Let 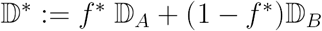 and 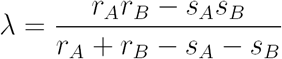. Then, 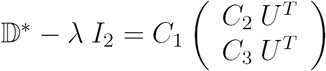, where 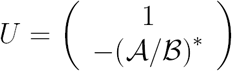 along with

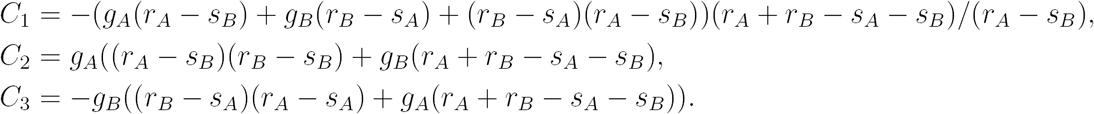

Since *U^T^V* =0 where *V = ((r_B_ – s_A_)=(r_A_ – s_B_), 1)^T^, (λ, V)* is an eigen pair of 픻*.

**Theorem A.10.** *In Stage 2 of the optimal strategy, both A_R_ and B_R_ change with a constant net-proliferation rate,*

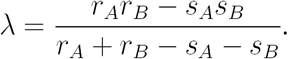

*Proof.* Without a loss of generality, let us prove it only when *A/B*(0) < (*A/B*)*.

If *A/B*(0) < (*A/B*)*, *DrugA* has a better effect initially. So following the optimal therapy scheduling, *DrugA* is assigned alone at the beginning as long as 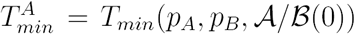 (Stage 1), and then Stage 2 starts at 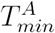 with initial condition

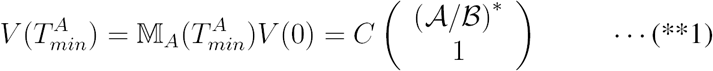

where 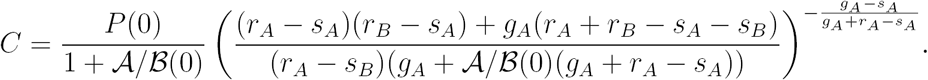

By Theorem A.8, in Stage 2, *V (t)* obeys

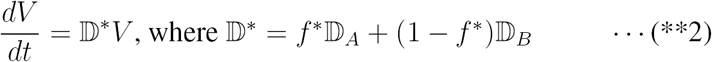

By Lemma A.9, *V* (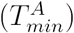) is an eigenvector of 픻* with the corresponding eigenvalue, *λ*. Then, the solution of (**2) with the initial value (**1) is

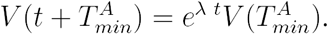

**Theorem A.11.** *With optimal therapy utilizing DrugA and DrugB, V obeys the following equations and solutions.*

If *A/B*(0) < (*A/B*)*,

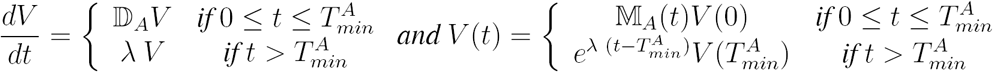

*Similarly if A/B*(0) ≥ (*A/B*)*,

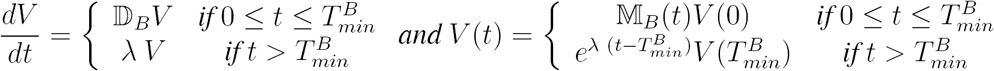

*Proof.* Straightforward, by Theorem A.10

## Appendix B Sensitivity analysis on optimal scheduling

The two determinant quantities of optimal control scheduling are (i) the duration of the first stage 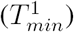, and (ii) the relative intensity between two drugs in the second stage (*k\**). Here, we show sensitivity analysis on the quantities related to them, *T_gap_* and *f*\*, over a range of (scaled) model parameters. Additionally over the same range, we studied how much our *T_min_-*based optimal scheme is better than the *T_max_-*based scheme evaluated by the integral in equation (13).

1. Sensitivity analysis of *T_gap_*
Using *g*_1_, we non-dimentionalize all the values, like

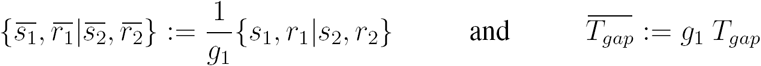

then,

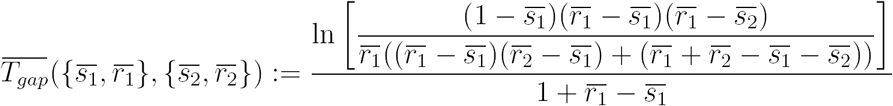

In general, cells mutate slower than they proliferate, so we ran sensitivity analysis on *T_gap_* for all *a* 1 for 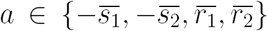. Figure 12 shows *T_gap_* over the range of 20 < 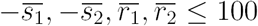. So, under the assumption that *g*_1_ min{–s_1_, –*s_2_, r_1_ r_2_}*,

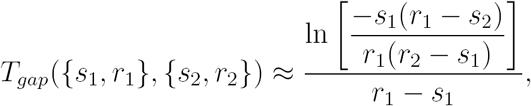

which approximates the contour curves of Figure 12.

**Figure 12:**
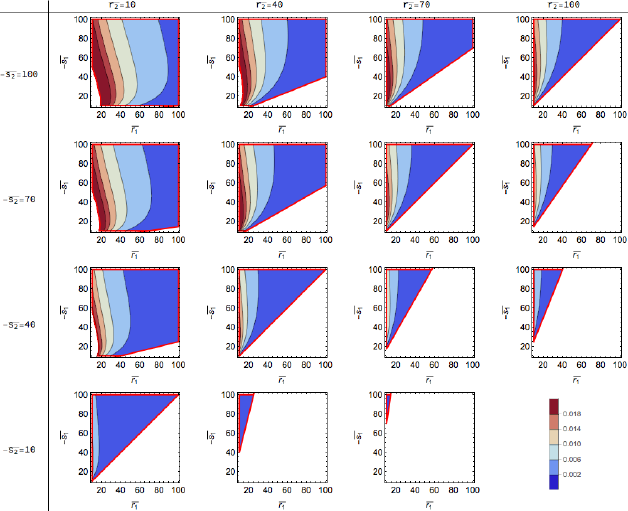
**Contour maps of** *T_gap_* over ranges of 10 ≤ *a* ≤ 100 for 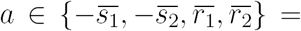 {–*s_1_,–s_2_, r_1_; r_2_*}/*g*1 and *r_1_r_2_* < *s_1_s_2_* (Condition (6)). As –*s*_2_ decreases and/or *r*_2_ increases, the optimal switching timing to the second drug is delayed (*T_min_* ↑ and *T_gap_* ↓). As *r*_1_ increases, Tgap decreases. Also, *T_gap_* and *s*_1_ have a non-monotonic relationship as shown on the graphs.

2. Sensitivity analysis of *f*\*

Regarding the regulated intensities among the two drugs, *k\**, we assumed that *g*_1_ ~ *g_2_* := *g*, similarly assuming that they are both much smaller than {– *s_1_, –s_2_, r_1_ r_2_*). Then we normalized all the parameters with the unit of g, like

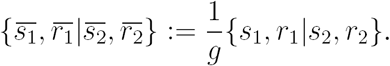

*k*\* can be rewritten in terms of the dimensionless parameters.

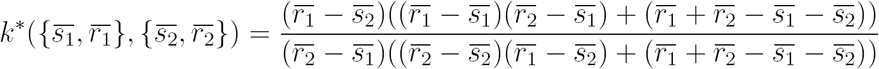

In this sensitivity analysis, we use

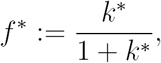

which represents intensity fraction of the initially better drug out of the total therapy. We evaluated *f*\* over the same ranges of *{s_1_, s_2_,r_1_,r_2_}*, like the previous exercise (see Figure 13) over the range max{*g_1_, g_2_*} ≪ min{–s_1_, – *s_2_, r_1_, r_2_}*, so *k\** and *f*\* can be approximated by the simpler forms:

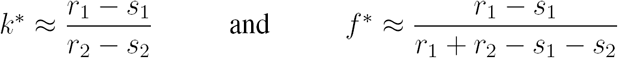

**Figure 13:**
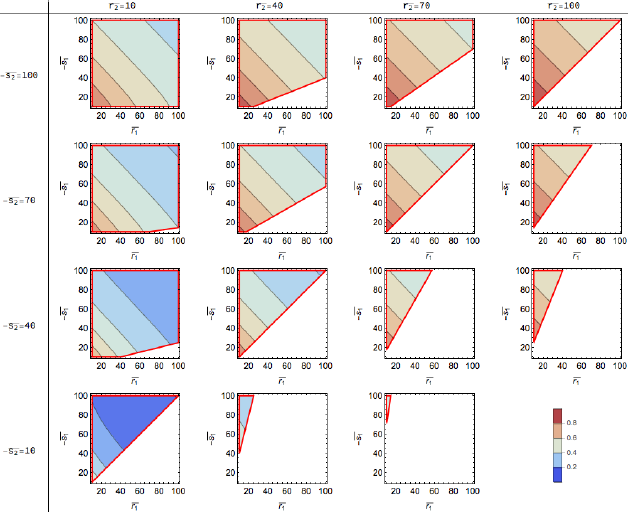
**Contour maps of** *f*\* over ranges of 10 ≤ *a* ≤ 100 for *a* 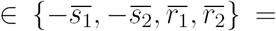 {–*s*_1_,–*s*_2_,*r*_1_,*r*_2_}/*g* and *r*_1_*r*_2_ < *s*_1_*s*_2_ (Condition 6). k* (or f*) increases, as *r*_1_ and/or –*s*_1_ decreases and/or as *r*_2_ and/or –_s_*2* increases.

3. Sensitivity analysis of Integral (13)

To study the sensitivity of the advantage of using the optimal control defined by Integral (13), we assumed that *g*_1_ ≈ *g*_2_ ≈ *g* = 0.001. Then similar to the previous studies, we explored the sensitivity of the normalized parameters in terms of *g*, that is:

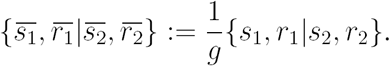

## Appendix C Clinical implementation of instantaneous switch in the optimal strategy

In clinical practice, the instantaneous drug-switch which we suggest in the second stage of the optimal treatment scheduling is not implementable. Therefore, we compared similar schedules to the optimal case. In the “similar” schedules, the first stage, using an initial drug, remained the same as the optimal schedule. However the second part, where we previously used an instantaneous switch (with Δ*t* = 0), was modified to use a fast switch (Δ*t* > 0). Figure 15 (a) and (b) shows how instantaneous switching (Δ*t* = 0) and fast switching (multiple choices of Δ*t* > 0) compare in terms of population size using different drug parameters. As expected, the smaller Δ*t* is, the closer to the ideal case. And, a choice of a reasonably small Δ*t* (like 1 day or 3 days) results in an outcome quite close to the optimal scenario.

We repeated this exercise with *k\** (from equation (12)) instead of *k(Δ*t**) modulated by Δ*t* (Figure 15 (c) and (d)). Only small differences are observed between Figure 15 (a) and (b) and Figure 15 (c) and (d), which justifies the general usefulness of *k\** independent of Δ*t*.

**Figure 14:**
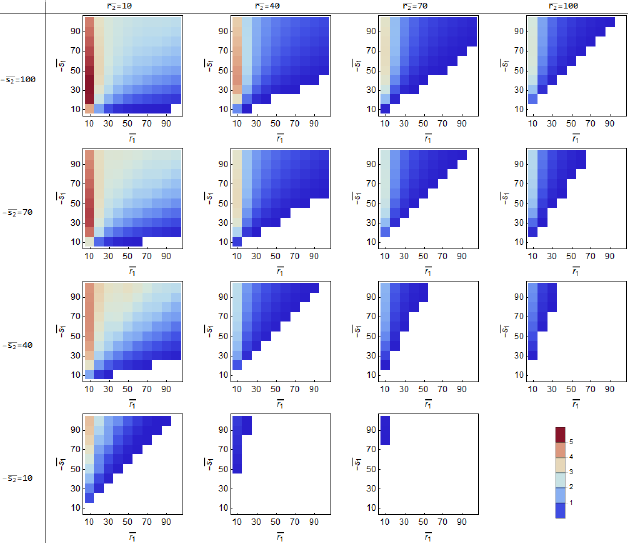
**Contour maps of the measured advantageous effect of the optimal therapy** defined by the integration (13) over ranges of 10 ≤ *a ≤* for 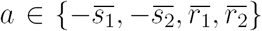 and *r*_1_*r*_2_ < *s*_1_*s*_2_(Condition (6)) Here, 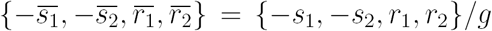 and *g* = 0.001. The measured effect increases as *r*_1_, *r*_2_ decreases and/or – *s*_1_ increases.

**Figure 15:**
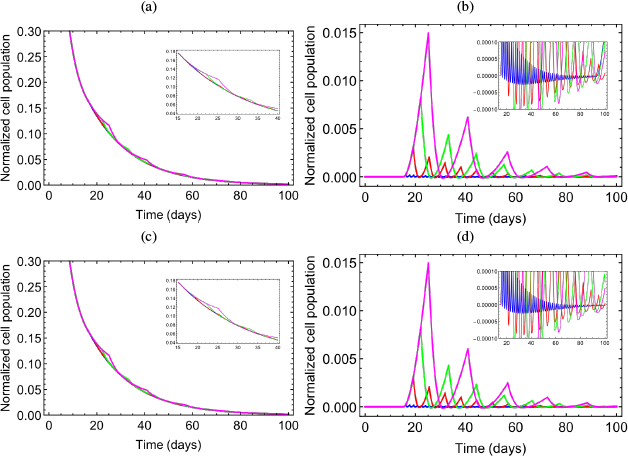
**Graphs showing regular drug switching in Stage 2 with different** *{Δ*t*, k{Δ*t*,p_A_,p_B_)}: Δ*t** =1 day (blue), *Δ*t** = 4 days (red), *Δ*t* =* days (green), and *Δ*t** = 10 days (magenta). Parameters/conditions: *p_A_* = {–0.18,0.008,0.00075}/day, *p_B_* = {–0.9,0.016,0.00125}/day and 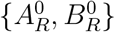 = {0.1,0.9} (a) Total population histories, 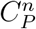 for *n* G {1,4,7,10} days (b) Differences between the optimal population history 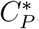, (i.e., when *Δ*t** = 0) and each case with positive *Δ*t**. (i.e., 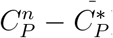). The inserts interesting ranges, (c) and (d) are equivalent with (a) and (b) except that *k*(p_A_,P_B_)}* has been used instead of *k(Δ*t*, p_A_, p_B_)}*

## Appendix D Stochastic simulation codes

The computational code written in Python will be provided at *Github* (https://github.com/nryoon12/Optimal-Therapy-Scheduling-Based-on-a-Pair-of-Collaterally-Sensitive-Drugs).

